# Nucleolar Dynamics During Oogenesis

**DOI:** 10.64898/2026.05.19.726235

**Authors:** Ruoyu Li, Grace McKown, Dai Tsuchiya, Mark Mattingly, Anna Galligos, Michay Diez, Jui Feng Lu, Mary C McKinney, Sean McKinney, Boris Rubinstein, Timothy J Corbin, Melainia McClain, Carrie Carmichael, Victoria A Hassebroek, Stephanie H Nowotarski, Jennifer L Gerton, Kamena K Kostova

## Abstract

Ribosome biogenesis is a conserved and highly regulated process that starts in the nucleolus, a membrane-less multi-phase organelle. Although the architecture of the nucleolus is known to change due to perturbations, how nucleolar organization is modulated during physiological processes to meet changing translational demands remains unclear. Here, we use zebrafish oogenesis as a developmental context requiring a rapid expansion of translational capacity to investigate the regulation of nucleolar architecture. We show nucleoli undergo coordinated changes in number, size, subnuclear localization, and layering throughout oogenesis. We further demonstrate that nucleoli form around extrachromosomal DNA circles that contain the rDNA locus. Notably, mouse oocytes undergo similar developmental changes in nucleolar layering and phase organization, indicating that remodeling of nucleolar condensates is a conserved feature of oogenesis. These findings reveal previously unexplored regulation of nucleolar architecture as developmental adaptations to changing biosynthetic needs.

## Introduction

Ribosome biogenesis, the assembly of mature ribosomal subunits, is a highly complex and conserved process involving 80 ribosomal proteins, 4 ribosomal RNAs (rRNAs), and over 200 processing factors^1^. This energy-intensive process must progress continuously in metabolically active cells to sustain protein synthesis and ensure cell survival^2^. Transcription of rRNAs and their incorporation into maturing ribosomal subunits occurs within nucleoli, liquid-like membrane-less organelles located in the nucleus^3,4^. Each nucleolus contains three nested liquid-liquid phase separated compartments: fibrillar center (FC), dense fibrillar component (DFC), and granular component (GC)^3,5,6^. rRNA synthesis occurs at the FC-DFC interface, while rRNA processing and ribosomal subunits assembly proceeds in the DFC and GC^5,7,8^.

In most somatic cells, nuclei contain a limited number of nucleoli, yet their number, morphology, and phase organization can change dramatically under stress^9,10^. For example, inhibition of RNA synthesis with Actinomycin D induces the formation of nucleolar caps in which FC and DFC phases re-localize outside of the GC^9^. Other perturbations, including cell cycle dysregulation and aging, are associated with nucleolar enlargement and fusion^11,12^. Together, these observations indicate that nucleolar architecture is dynamically remodeled in response to cellular demand. However, how this architecture is regulated during normal physiological processes to meet changing translational demand remains poorly understood.

Addressing this question requires examining a developmental trajectory in which protein synthesis demands are tightly orchestrated. Oogenesis provides such a system, as the growing oocyte undergoes coordinated increases in size and biosynthetic capacity. Electron microscopy studies in *Xenopus* and bovine oocytes suggest that the nucleolar architecture is extensively remodeled during oogenesis^13–15^. For example, *Xenopus* oocytes assemble thousands of nucleoli around amplified copies of the rDNA loci^16,17^. More recent studies of late-stage mouse^18–20^ and *Xenopus* oocytes^6,13,21^ have provided insights into nucleolar organization, and the biophysical principles governing the behavior of this liquid-like organelle^5^. However, how the nucleolar architecture is established and maintained throughout the developmental trajectory remains poorly understood.

*Danio rerio* provides a powerful system to address this question. Zebrafish produce large numbers of oocytes on a weekly basis, and oogenesis is asynchronous, yielding follicles at all stages of development within a single ovary (Fig. 1a)^22^. Over approximately two weeks, oocytes grow from approximately 50 µm to 800 µm in diameter, resulting in a 3,500-fold increase in volume^22^. This extraordinary expansion requires a dramatic increase in ribosome biogenesis and protein synthesis capacity. Importantly, zebrafish exhibit developmentally regulated ribosome heterogeneity: maturing oocytes contain maternal ribosomes, whereas somatic cells contain zygotic ribosomes. These two ribosome types differ in all four rRNAs and are encoded by distinct genomic loci^23,24^. The sequence differences between maternal and zygotic rRNA enable selective detection of oocyte-derived rRNA and provide a unique opportunity to directly monitor maternal ribosome biogenesis *in vivo*. This is particularly important because oocytes are surrounded by biosynthetically active support cells that can synthesize and transfer biomolecules into the developing oocyte^25,26^, making it challenging to distinguish cell-autonomous from non-autonomous ribosome production in most systems.

**Fig. 1:**
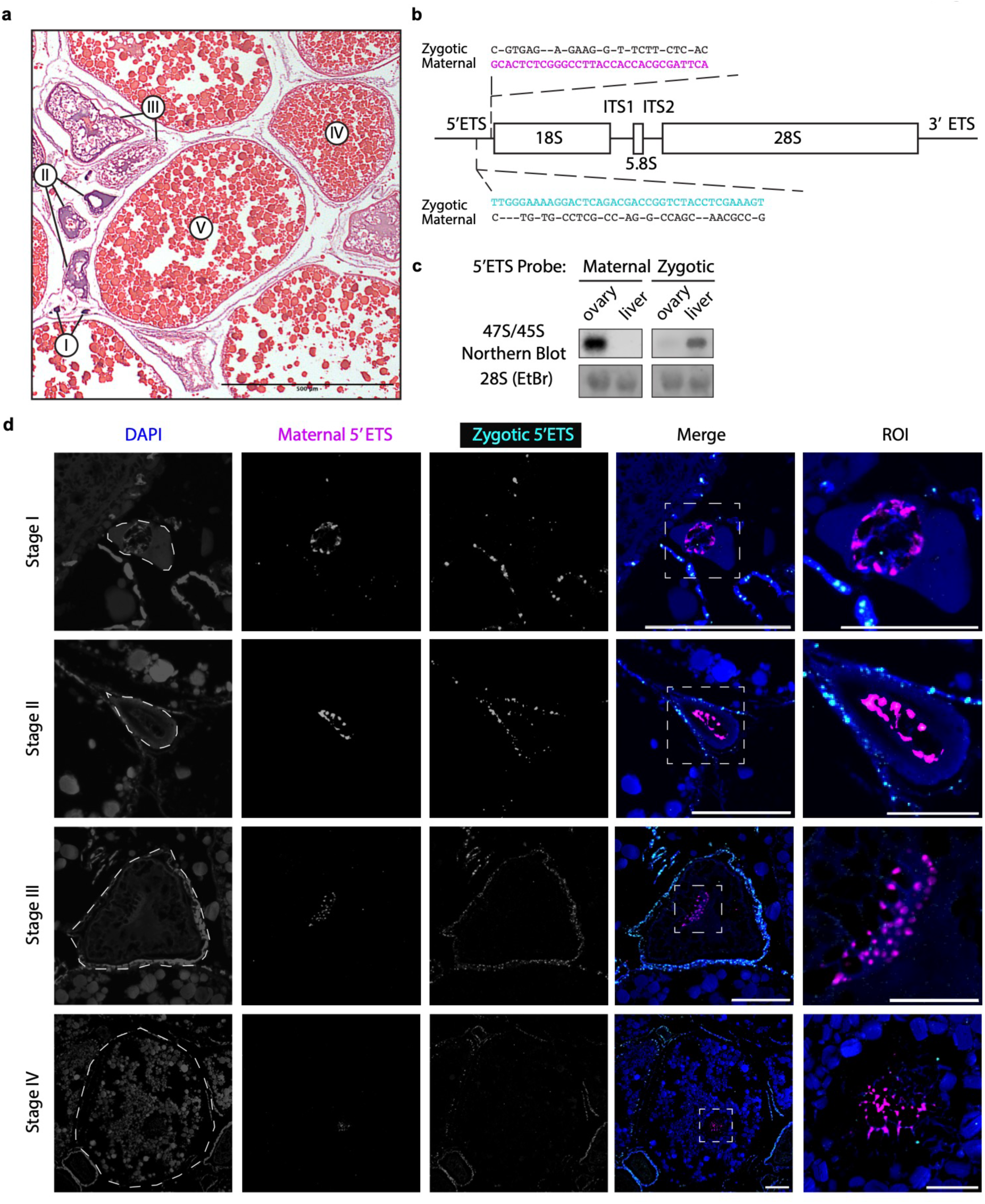
Maternal ribosome production in *Danio rerio* occurs exclusively within oocytes. (**a**) Hematoxylin and Eosin (H&E) staining of paraffin sections from zebrafish ovary. Highlighted are examples of stages I-V oocytes. Scale bar: 500 µm. (**b**) Diagram of the 45S ribosomal RNA precursor indicating the 5′ and 3’ external transcribed spacers (5′-ETS and 3’ETS), internal transcribed spacers (ITS1 and ITS2), and mature 18S, 5.8S, and 28S rRNA regions. Probes targeting maternal (magenta) and zygotic (cyan) 5′ETS are shown, with the corresponding zygotic or maternal sequence highlighted in black. Note that the presence of mutations and indels ensures that each probe specifically recognizes its respective locus without cross-reactivity. (**c**) Northern blot analysis of nascent rRNA isolated from ovary or liver. Probes highlighted in A are used to detect the maternal and zygotic 45S rRNA. Ethidium bromide (EtBr) staining of mature 28S ribosomal RNA is serving as a loading control. (**d**) HCR RNA-FISH of 5’ETS maternal (magenta) and zygotic (cyan) rRNA in paraffin-sectioned oocyte stages I-IV. Approximate oocyte border indicated in white dashed line. Nuclei are stained with DAPI (blue). Region of interest (ROI) highlights the maternal 5’ETS signal in the nucleus of oocytes. Scale bars: 100 µm in first four columns, 40 µm in last column.

In this study, we show that rRNA synthesis is cell-autonomous during zebrafish oogenesis. We find that nucleolar architecture undergoes coordinated changes in number, morphology, volume, subnuclear localization, and FC–DFC–GC organization throughout follicle development. These architectural transitions are supported by nuclear F-actin, as disruption of F-actin assembly leads to nucleolar coalescence. We further demonstrate that nucleoli form around extrachromosomal rDNA amplified from the maternal rDNA locus. Notably, mouse oocytes undergo similar developmental remodeling of nucleolar layering and phase organization, indicating that dynamic regulation of nucleolar architecture is conserved across species. Together, these findings establish oogenesis as a developmental context in which nucleolar architecture is actively remodeled to meet changing translational demands, revealing regulatory principles inaccessible in conventional reductionist systems, such as cell lines or unicellular organisms.

## Results

### Autonomous ribosome production during oogenesis in zebrafish

Whether ribosome production is autonomous during zebrafish oogenesis remains unknown. In zebrafish, the mature oocyte contains a distinct type of ribosomes, termed maternal ribosomes, whereas somatic tissues examined to date assemble zygotic ribosomes^23,24^. These ribosomes differ in both the mature rRNA sequences (5S, 5.8S, 18S, and 28S rRNA), as well as the transcribed spacer sequences (5’ external transcribed spacer (5’ETS), internal transcribed spacer 1 (ITS1), internal transcribed spacer 2 (ITS2), and 3’ external transcribed spacer (3’ETS)) that are removed during ribosome biogenesis^23^. These sequence differences provide a means to identify where in the ovary maternal rRNA is synthesized.

We designed hybridization chain reaction fluorescence *in situ* hybridization (HCR-FISH^27,28^) probes that target the maternal or zygotic 5’ ETS (Fig 1b), a region of the rRNA that is rapidly processed during ribosome biogenesis^29^, and as a result is only found in the nucleolus and not in the cytoplasm as part of the mature ribosome. Northern blot analysis confirmed that the maternal and zygotic 5’ETS probes detected ribosome biogenesis intermediates in ovary and liver respectively (Fig. 1c). The faint signal observed in the ovary sample using the zygotic 5’ETS probe is due to the presence of somatic tissue, such as muscle, connective tissue and granulosa cells in the ovary^30^.

HCR-FISH staining of paraffin sections from zebrafish ovaries, using probes specific to maternal and zygotic nascent rRNA, revealed strong maternal signal within oocytes, while surrounding granulosa and theca cells exhibited zygotic signal (Fig. 1d, Fig. S1). This pattern was consistent across oocyte stages I–IV. Together, these observations demonstrate that maternal rRNA synthesis occurs exclusively within the oocyte and is autonomous throughout zebrafish oogenesis.

### Dynamic changes in nucleolar numbers, shapes, and localization during oogenesis

Unexpectedly, oocytes at distinct stages of oogenesis exhibit marked differences in nucleolar number, morphology, and localization (Fig. 1d). However, using paraffin sectioning to examine the architecture of the nucleoli is challenging due to the large size of the maturing oocytes and the potential for tissue distortions during fixation and processing that disrupt overall morphology. Furthermore, zebrafish oocytes are lipid-rich, which causes high autofluorescence. To address these limitations, we optimized tissue clarification protocol in combination with HCR-FISH staining and imaging of whole-mount zebrafish oocytes. We used 5’ETS HCR-FISH to define nucleoli. This new method allowed us to perform 3D reconstructions of the nucleoli and gain a comprehensive understanding of the nucleolar dynamics during oogenesis (Fig. 2a).

**Fig. 2:**
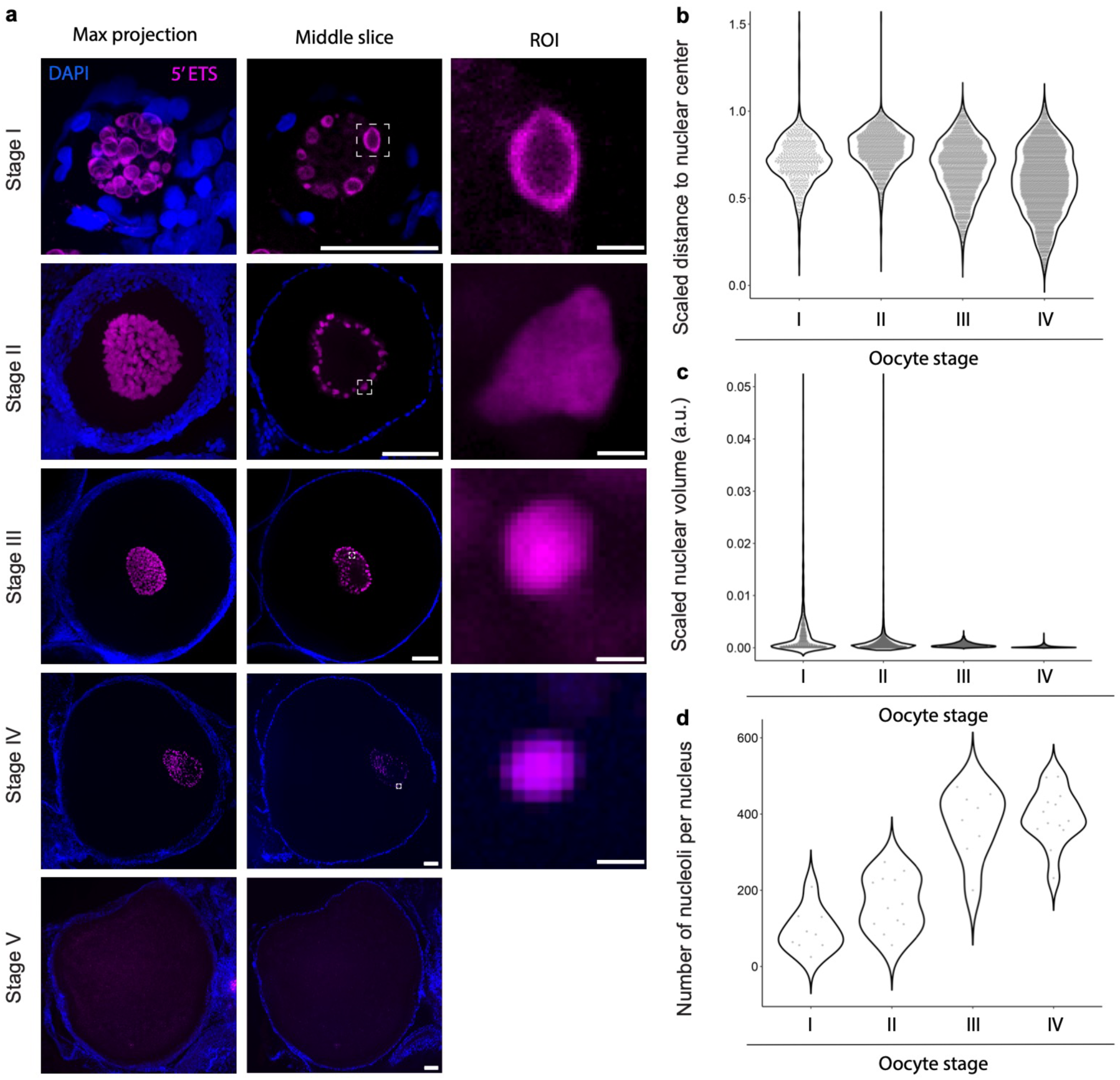
Nucleolar morphology is highly dynamic throughout oogenesis. (**a**) HCR RNA-FISH of 5’ETS maternal rRNA (magenta) in whole-mount oocyte stages I-V. Nuclei are stained with DAPI (blue). Individual nucleoli are highlighted in the region of interest (ROI) indicated in dotted white box in middle column. Scale bar: stage I, 50 µm; stages II-V, 100 µm; individual nucleoli highlighted in last column, 5 µm. (**b**) The subnuclear localization of each nucleolus was measured by comparing the distance between the centroid of each nucleolus to the centroid of the nucleus. Nucleoli close to the nuclear periphery have a scaled distance close to 1, whereas nucleoli close to the center of the nucleus have a scaled distance close to 0. Points outside the displayed y-axis range are not shown but were retained in the violin density calculation. Metrics are defined quantitatively in Materials and Methods section. (**c**) Quantification of nucleolar volume, scaled to nuclear volume in oocyte stages I-IV. Points outside the displayed y-axis range are not shown but were retained in the violin density calculation. (**d**) Quantification of number of nucleoli per oocyte nucleus in stages I-IV. (**b-d**) Stage I, N=8 oocytes, 848 nucleoli; Stage II, N=13 oocytes, 2003 nucleoli; Stage III, N=8 oocytes, 3012 nucleoli; Stage IV, N=13 oocytes, 5087 nucleoli.

We discovered stage-specific changes in the nucleolar number, shape, volume, and sub-nuclear localization (Fig. 2b-d). Stage I oocytes contain approximately 100 nucleoli that vary widely in size. In most nucleoli examined, the 5′ETS signal is confined to the periphery, whereas the center is depleted of signal, resulting in a soap bubble-like architecture (Video S1). Furthermore, the nucleoli are localized to the nuclear periphery. Similar to stage I nucleoli, nucleoli in stage II oocytes are also localized to the nuclear periphery (Video S2). However, these nucleoli have irregular shapes and more uniform sizes compared to stage I nucleoli. The number of nucleoli also increased between stage I and stage II oocytes. These data suggest either *de novo* assembly of nucleoli or nucleolar fission events, where a pre-existing nucleolus divides to give rise to two new nucleoli. Both mechanisms can potentially increase ribosome biogenesis capacity during oocyte maturation. During stages III and IV, the nucleoli adopt a round shape with uniform HCR signal, similar to the signal observed in somatic cells. Importantly, these nucleoli change sub-nuclear localization and migrate towards the center of the nucleus. Although the number of nucleoli increases between stage II and III oocytes, the average number of nucleoli does not change in stage IV oocytes (Video S3, S4). Finally, in stage V oocytes the nuclear membrane is dissolved, and the oocyte is ready for fertilization. Consistent with previous literature, rRNA synthesis is inhibited in mature oocytes^31^, and no 5’ETS HCR-FISH signal is detectable.

### Nuclear actin polymerization is required to maintain nucleolar number

Work in late-stage *Xenopus* oocytes and early *Drosophila* oocytes revealed that nuclear filamentous actin (F-actin) stabilizes nucleolar structure^15,24,25^. In *Xenopus*, depolymerization of the oocyte nucleoskeleton results in coalescence of the nucleoli^6,24^. To investigate if F-actin plays a role in maintaining nucleolar architecture throughout zebrafish oogenesis, we treated oocytes (stages I–IV) with Latrunculin A (Lat-A) (Fig. 3a), a macrolide that disrupts F-actin formation by sequestering actin monomers^26^. Phalloidin staining, which specifically labels F-actin, revealed a dense nuclear network at all stages. This network appeared more fibrous in stage I–II oocytes and formed a dense meshwork in stage III–IV oocytes. Notably, this network contained discrete gaps that corresponded to nucleolar regions, consistent with nucleoli being embedded within, and excluded from, the surrounding F-actin matrix.

**Fig. 3:**
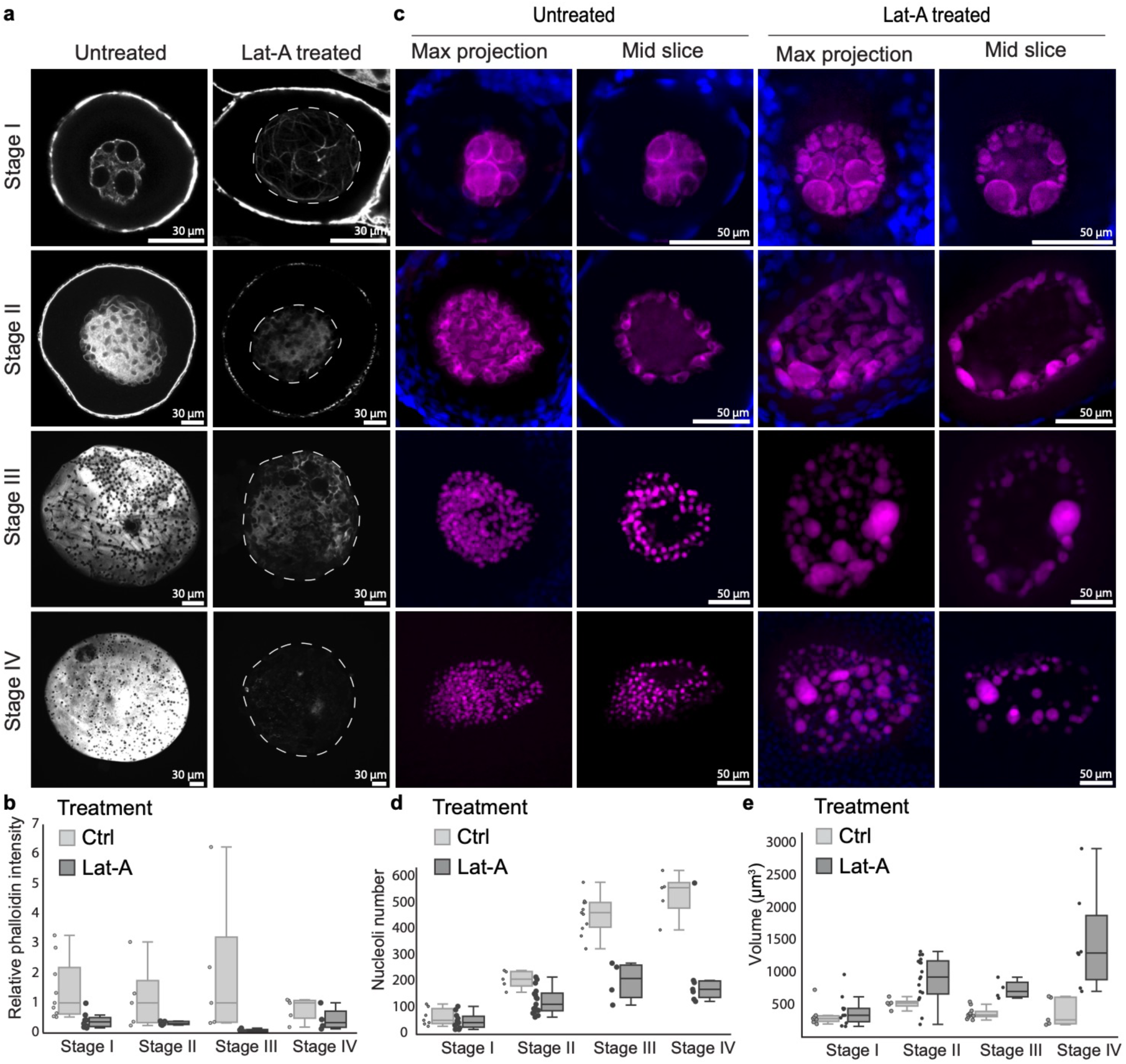
A nuclear F-actin matrix maintains nucleolar morphology. (**a,c**) Phalloidin staining of F-actin (a) or HCR RNA-FISH for maternal 5’ ETS (b) in whole-mount control oocytes or in oocytes treated with Latrunculin A (Lat-A). Scale bar: 30 µm (a) and 50 µm (b) (**b**) Relative intensity of Phalloidin staining in control oocytes or oocytes treated with Lat-A. For control group, Stage I, N=9; Stage II, N=5; Stage III, N=5; Stage IV, N=5. For Lat-A treated group, Stage I, N=8; Stage II, N=6; Stage III, N=5; Stage IV, N=4. (**d,e**) Quantitation of number (d) and volume in µm^3^ (e) of nucleoli in individual oocytes with and without Lat-A treatment. For control group, Stage I, N=7; Stage II, N=7; Stage III, N=11; Stage IV, N=8. For Lat-A treated group, Stage I, N=11; Stage II, N=17; Stage III, N=8; Stage IV, N=7.

Lat-A treatment disrupted nuclear F-actin structure across all four stages (Fig. 3a, b). To determine how this disruption impacts nucleolar architecture, we performed HCR-FISH using probes against the maternal 5′ETS (Fig. 3c). Disruption of F-actin reduced the number of nucleoli per cell (Fig. 3d) and promoted the formation of larger nucleoli across all oocyte stages (Fig. 3c, e), consistent with nucleolar coalescence. This effect was most pronounced in stage III–IV oocytes, where multiple nucleoli frequently merged into substantially larger structures compared to untreated controls.

Together, these data indicate that F-actin constrains nucleolar coalescence throughout oogenesis. Unexpectedly, disruption of F-actin did not alter subnuclear positioning: stage I–II nucleoli remained peripheral, whereas stage III–IV nucleoli remained distributed throughout the nuclear volume after Lat-A treatment (Fig. 3b). These findings suggest that additional structural components govern nucleolar positioning during oogenesis.

### Nucleolar layering is established late in oogenesis

HCR-FISH imaging of the maternal 5’ETS revealed that nucleolar architecture is dynamically regulated during zebrafish oogenesis (Fig. 2). The nucleolus is a membraneless, multiphase organelle with three spatially organized and nested layers, two of which are defined by the presence of 5′ETS-containing nascent rRNA (Fig. 4a). To further investigate how nucleolar phases change throughout oogenesis, we used immunostaining of FC and DFC/GC markers (Fig. 4b, c). Ubtf is a transcription factor located in the FC that facilitates Pol I-rDNA promoter interaction and subsequent rRNA synthesis. Nucleolin (Ncl), a DFC/GC marker, binds to pre-rRNA and small nucleolar ribonucleoprotein (snoRNP) complexes to mediate rRNA processing^28^. Under the canonical model of nucleolar organization, Ubtf localizes to discrete puncta that reside within the Ncl-defined compartment. Surprisingly, Ubtf and Ncl showed co-localization in stage I and II oocytes, indicating a lack of canonical FC–DFC/GC phase separation. This colocalization was observed across nuclei of varying sizes and independent of their subnuclear localization, suggesting that early oocyte nucleoli do not have canonical layering (Fig. S2). In contrast, Ubtf puncta positioned interior to the Ncl signal were detected in stage III and IV oocytes (Fig. 4 and S2), indicating that canonical FC–DFC/GC organization is established later during oogenesis.

**Fig. 4:**
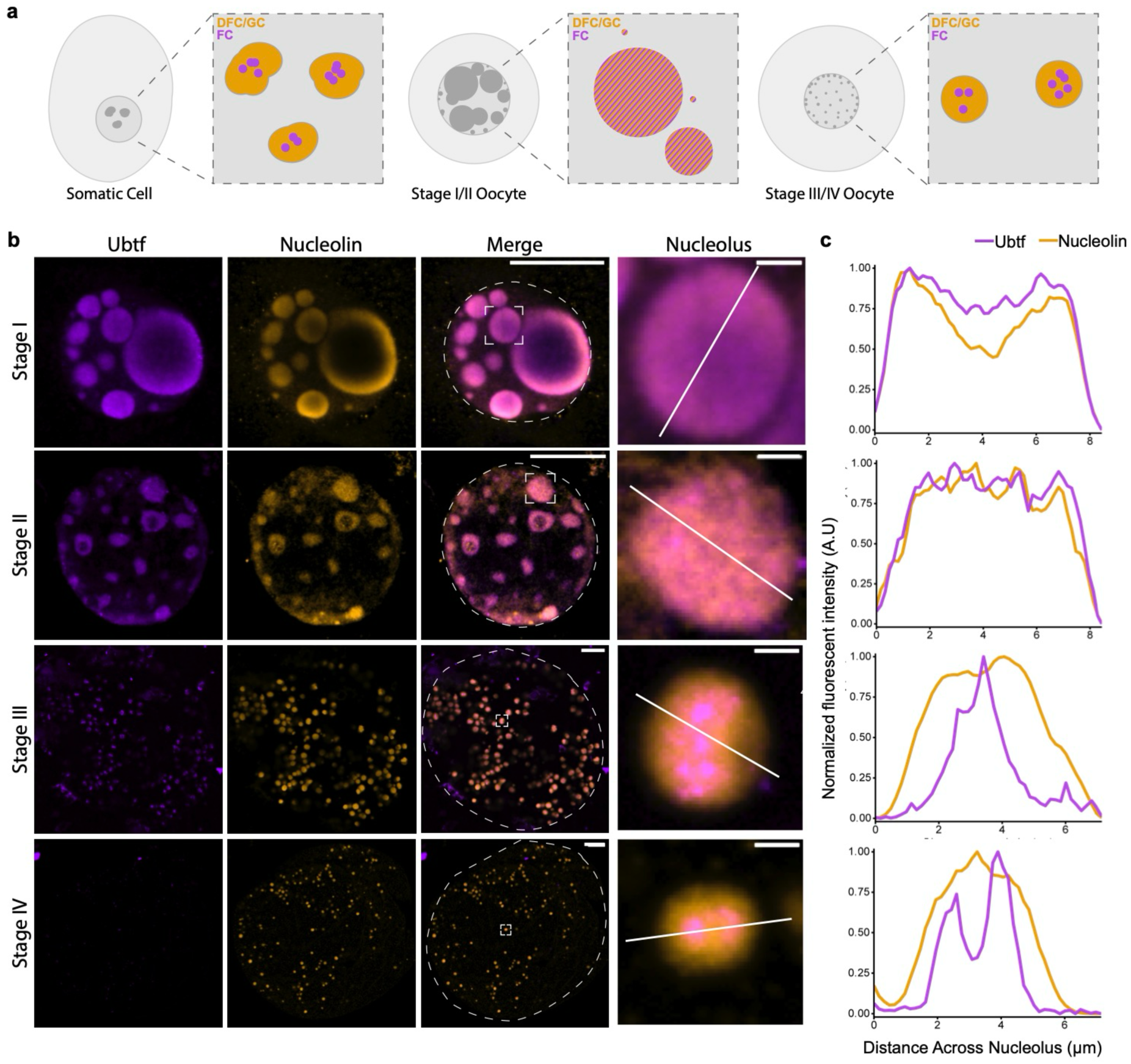
Nucleolar layering is not established until mid-late oogenesis. (**a**) Schematic of nucleolar architecture in somatic cells and oocytes. Fibrillar center (FC) is colored in purple, dense fibrillar component and granular component (DFC/GC) are colored in orange, and nucleoplasm is colored in gray. (**b**) Immunostaining of Ubtf (purple) and Nucleolin (orange) in whole-mount stage I-II oocytes and mounted nuclei of stage III-IV oocytes. A single optical z-slice through is shown. Brightness and contrast were adjusted for visualization. Approximate nuclear border is indicated with white ellipse. A single nucleolus (white box) was selected for fluorescent intensity measurements indicated by white line. Scale bars: merge, 20 µm; nucleolus: 2 µm. (**c**) Quantitation of fluorescent intensities of Ubtf and Nucleolin across nucleolus shown in (b). Intensities were normalized to minimum and maximum values. A. U., Arbitrary Units.

It has been postulated that only nucleoli in amniotes have a tripartite nucleolus, whereas other organisms, including zebrafish, assemble bipartite nucleoli that contain a fibrillar zone (FZ) and granular zone (GZ)^36,37^. Determining whether the Ubtf puncta nested in the Ncl signal in the nucleoli of stage III and IV oocytes represent a true FC or FZ is challenging due to the limited number of antibodies for zebrafish samples. Instead, we used electron microscopy to explore in further detail if late-stage oocytes contain three distinct nucleolar layers (Fig. S3). Indeed, our nucleolar reconstructions from stage IV oocytes showed clear FC-DFC-GC compartments. These observations raise the possibility that tripartite nucleoli are more ubiquitous than previously expected, and their formation might correlate with the translational needs of the cell rather than the taxonomy of the species.

Together, these results indicate that nucleolar layering can be altered not only due to perturbations of rRNA synthesis^7,9^, but also as part of normal cellular maturation, such as oogenesis, where the nucleoli are initially unstructured and undergo progressive establishment of canonical FC–DFC–GC phase organization during maturation.

### Maternal rDNA is amplified, circularized, and used as a template for rRNA synthesis

Oocytes face a unique challenge in meeting rRNA demand: they must produce vast quantities of rRNA from a fixed number of rDNA loci while maintaining genomic integrity, precluding polyploidy or polyteny. Many organisms address this constraint by selectively amplifying rDNA and depositing these extrachromosomal copies in the oocyte^38–40^. Zebrafish contains a single maternal rDNA locus on chromosome 4^23^, yet it can assemble hundreds of nucleoli that actively synthesize rRNA (Fig.2d). To test whether these nucleoli are organized around extrachromosomal DNA (ec DNA), we performed fluorescent *in situ* hybridization (FISH) with rDNA probes in whole-mount zebrafish oocytes (Fig. 5a). Extrachromosomal rDNA (ec rDNA) was detected at all oocyte stages. Notably, its subnuclear localization mirrored that of nucleoli, defined by 5’ETS HCR-FISH signal, with ec rDNA positioned near the nuclear periphery in stage I–II oocytes and distributed throughout the nucleoplasm in stage III–IV oocytes.

**Fig. 5:**
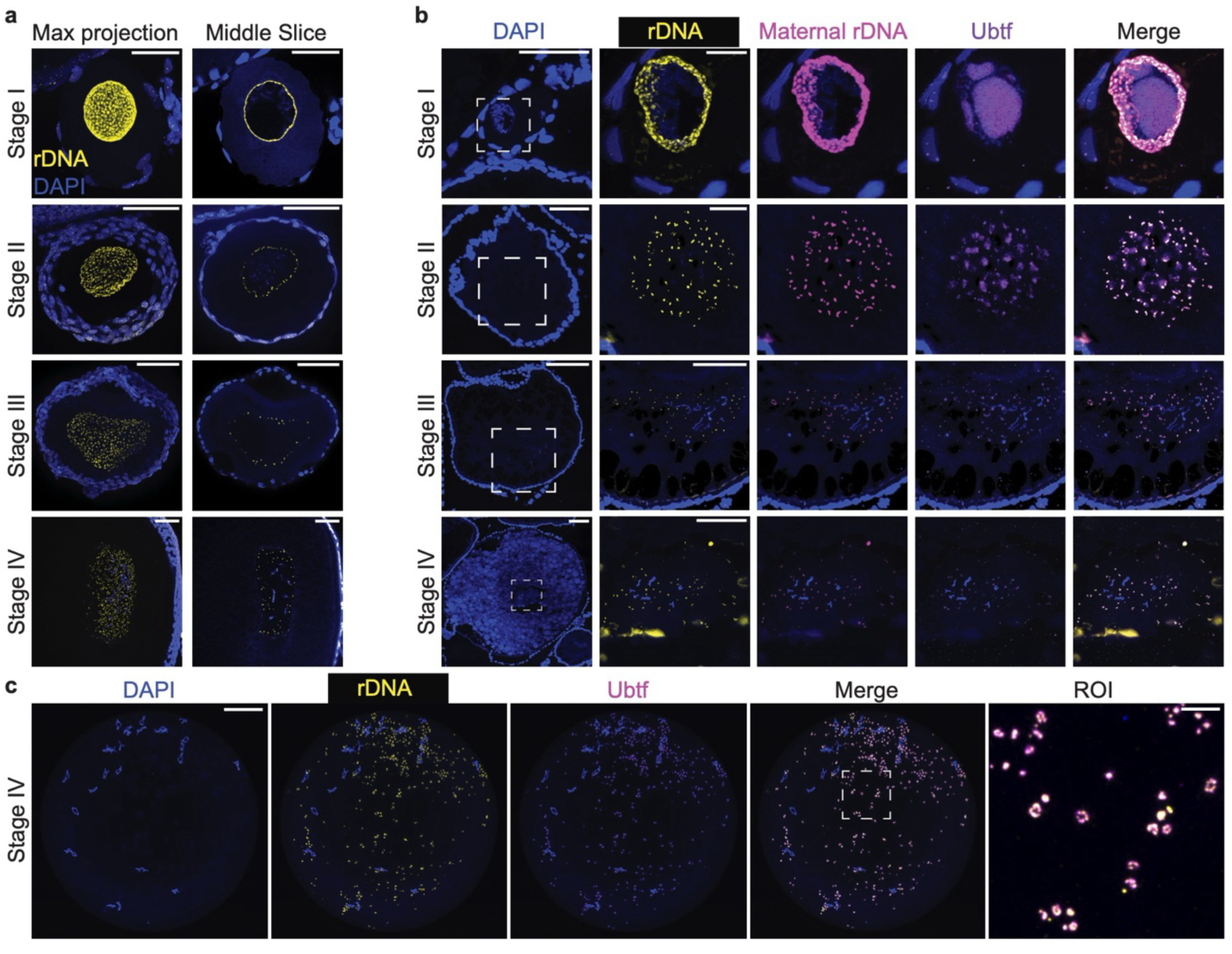
rDNA is amplified and deposited as extrachromosomal circles during zebrafish oogenesis. **(a)** rDNA FISH on whole oocytes (stages I-IV) embedded in hydrogel showing rDNA in yellow, and DAPI in blue. Brightness and contrast were adjusted for visualization. Scale bars: stage I, 50 µm; stages II-IV, 100 µm **(b)** Fluorescence in situ hybridization (FISH) of zebrafish ovary cryosections containing oocytes at stages I–IV, using a universal rDNA probe (yellow) and a maternal intergenic spacer (IGS) probe (magenta), combined with immunofluorescence for UBTF (purple). Region of interest (ROI) of oocyte nucleus depicted in white box. Brightness and contrast were adjusted for visualization. Scale bars: stage I (DAPI, 50 µm; ROI, 10 µm), stage II (DAPI, 50 µm; ROI, 20 µm), stages III/IV (DAPI, 100 µm; ROI, 50 µm). **(c)** Isolated stage IV oocyte nucleus showing colocalization of extrachromosomal rDNA (ec rDNA; yellow), detected by FISH, and UBTF (purple), detected by immunofluorescence. ROI shown in right-most image depicted in white box. Brightness and contrast were adjusted for visualization. Scale bar: nucleus, 50 µm; ROI, 10 µm.

To determine the origin of this ec rDNA, we performed double FISH using universal rDNA probes and probes specific for the maternal intergenic spacer (IGS) (Fig. 5b). All ec rDNA puncta were double-positive, indicating that they are derived from the maternal rDNA locus. We next asked whether all ec rDNA copies are transcriptionally active. We assessed the activity of individual loci using Ubtf, a marker of active rDNA transcription^41,42^. In stage I oocytes, only a subset of ec rDNA puncta colocalized with Ubtf, which instead formed large structures in the nucleoplasm, indicating that many amplified loci are initially inactive. In stage II oocytes, all ec rDNA puncta colocalized with Ubtf, although individual Ubtf domains often encompassed multiple ec rDNA foci, suggesting shared transcriptional hubs. By stage III–IV, each ec rDNA punctum was associated with Ubtf, indicating widespread activation.

To further resolve the organization of ec rDNA in late-stage oocytes, we performed FISH combined with Ubtf immunofluorescence on isolated stage IV nuclei (Fig. 5c). High resolution imaging revealed ring-like ec rDNA structures, consistent with circular DNA. These structures contained multiple discrete Ubtf puncta, suggesting that each ecDNA molecule harbors multiple rDNA repeats capable of supporting transcription. These results indicate that zebrafish oocytes amplify the maternal rDNA locus into extrachromosomal arrays that are spatially organized, developmentally activated, and structurally configured to support the high levels of rRNA synthesis required during oogenesis.

### Changes in nucleolar layering are conserved during M. musculus oogenesis

Since nucleoli undergo regulated changes in number, size, subnuclear localization, and layering during zebrafish oogenesis, we wanted to explore whether these dynamics are conserved in other vertebrates, including *M. musculus*. Like in zebrafish, the sexually mature mouse ovary contains oocytes at all stages of development, from a 10 µm primordial follicle to an 80 µm early antral follicle before ovulation (Fig. 6a). Investigations of mouse oocyte nucleoli focus predominantly on mature oocytes^18,20,31^, and therefore miss the preceding developmental trajectory. Understanding this nucleolar program throughout oogenesis in *M. musculus* offers a direct path to understanding whether the observed nucleolar regulation is a fundamental feature of vertebrate oocyte biology.

**Fig. 6:**
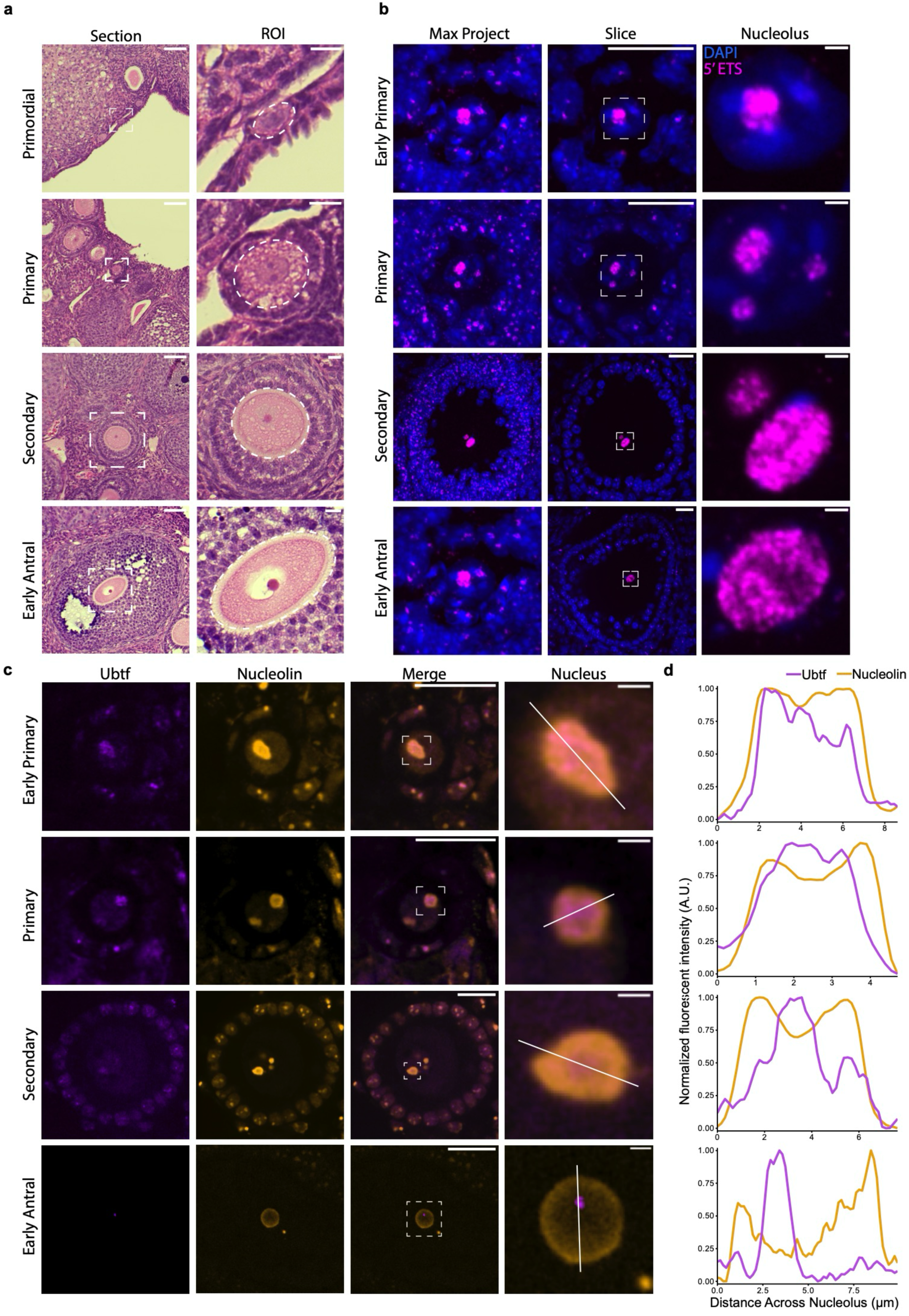
Dynamic regulation of nucleolar volume and layering in mouse oocytes. (**a**) H&E staining of mouse ovary paraffin sections with example oocytes of each stage. Approximate oocyte outlined in white ellipse. Scale bar: 20 µm. (**b**) HCR RNA-FISH for 5’ETS in whole-mount early primary, late primary, secondary and early antral oocytes. rRNA is shown in magenta and DAPI is shown in blue. Region of interest (ROI) indicated in dotted white box in middle column. Scale bars: slice, 20 µm, nucleolus, 2 µm. (**c**) Immunostaining of UBF (purple) and NCL (orange) in whole-mount oocyte stages early primary-early antral. Brightness and contrast were adjusted for visualization. A single nucleolus (white box) was selected for fluorescent intensity measurements indicated by white line. Scale bars: merge, 20 µm; nucleus: 2 µm. (**d**) Quantitation of fluorescent intensities of UBF and NCL across nucleoli shown in (d). Intensities were normalized to minimum and maximum values. A. U., Arbitrary Units.

To investigate nucleolar morphology throughout murine oogenesis, we visualized nucleoli using HCR RNA-FISH for mouse 5’ETS rRNA. Since primordial and late antral follicles exhibit minimal transcription and rRNA synthesis directly impacts nucleolar morphology, we focused our analysis only on stages that actively assemble ribosomes. Unlike zebrafish, mice express a single type of rRNA shared by all cell types, with no oocyte-specific isoform. As a result, the 5’ETS HCR-FISH probe labeled nucleoli in all oocytes and surrounding granulosa and theca cells (Fig. 6b). rDNA amplification has not been documented in mouse oocytes, and therefore nucleolar number does not increase throughout oogenesis. We performed immunostaining for upstream binding transcription factor (UBF) and nucleolin (NCL) to examine nucleolar phase layering (Fig. 6d, e, S4). Similar to zebrafish, mouse oocytes also exhibited stage-specific changes in their nucleolar layers. Early and late primary oocytes have overlapping UBF and NCL signals, whereas secondary oocytes exhibited the canonical UBF signal nested in NCL. In early antral follicles, nucleoli display a unique organization characterized by a ring-like NCL signal and a single UBF punctum positioned asymmetrically along the inner periphery of this ring.

These observations indicate that nucleolar architecture is dynamically regulated during oogenesis and exhibits both conserved and species-specific features. While we observe differences in nucleolar organization between zebrafish and mouse, likely reflecting differences in oocyte size and developmental timing, both systems show a dynamic regulation of the FC–DFC-GC phases during mid-to-late oogenesis. These shared features point to a conserved regulatory program that scales nucleolar organization to meet the increasing demand for ribosome production and to prepare the oocyte for early embryonic development, which relies on maternally deposited ribosomes. Given the many conserved aspects of oogenesis between mouse and human, these findings raise the possibility that similar regulation of nucleolar architecture occurs during human oogenesis.

## Discussion

Cells differ widely in their demands for protein synthesis and ribosome biogenesis, and growing oocytes represent an extreme case, requiring a massive expansion of translational capacity over a short developmental window. The mechanism by which the developing oocyte upregulates canonical ribosome biogenesis pathways or adopts new strategies to meet the protein synthesis demand is unclear. Here, we focus on the nucleolus, the site of rRNA synthesis and processing, and show that its organization is dynamically regulated during oogenesis. Specifically, we observe coordinated changes in nucleolar number, shape, subnuclear localization, and nucleolar layering. These findings indicate that nucleolar structure is actively remodeled to meet the increasing biosynthetic demands of the developing oocyte.

Synthesis of rRNA is a well-known rate limiting step of ribosome biogenesis^43^. To increase rRNA synthesis, zebrafish oocytes amplify the maternal rDNA locus and deposit the resulting DNA molecules as ec rDNA. These ec rDNA circles then serve as numerous nucleolar organizer regions (NORs). However, the mechanisms governing rDNA amplification, ecDNA activation, and turnover remain poorly understood. In *Xenopus*, rDNA amplification occurs during early oogenesis^38,39^ and has been proposed to proceed via rolling circle amplification^44,45^, with DNA polymerase alpha playing a critical role^46^. Our data show that zebrafish ec rDNA structures vary in size and in the number of associated Ubtf puncta (Fig. 5c). These observations suggest that each circle might contain a variable number of rDNA repeats, consistent with rolling circle amplification. Defining the factors that are involved in ec rDNA amplification and turnover and how they are regulated during oogenesis will be essential to understand how rDNA copy number and activity are regulated.

Ec DNA has recently emerged as a major driver of oncogene amplification in human cancers, where circular DNA molecules encoding growth-promoting genes can reach extremely high copy number, exhibit enhanced transcriptional activity, and segregate unevenly to fuel tumor evolution^47–49^. Although ecDNA formation in cancer is typically viewed as a pathological consequence of genome instability, it shares key architectural and functional features with developmentally programmed DNA amplification events, such as the formation of ec rDNA during oogenesis. These parallels raise the intriguing possibility that cancer cells co-opt or “hijack” latent cellular mechanisms for DNA amplification and extrachromosomal genome organization. Studying rDNA amplification during oogenesis therefore provides a tractable and physiologically relevant model to uncover the molecular logic of ec DNA formation, maintenance, and function, with direct implications for understanding how these processes are dysregulated in cancer.

A key feature of nucleolar regulation during oogenesis is the increase of the number of nucleoli, specifically between stages I and III (Fig. 2c). However, the mechanisms underlying this expansion are not clear. Changes in nucleolar number have been described in other contexts, such as mitosis and early embryogenesis. During mitosis in mouse and human cell lines, the nucleolus disassembles, but nucleolar organizing regions (NORs) persist as privileged chromosomal sites, with factors such as UBF remaining associated. At mitotic exit, processing factors first accumulate in prenucleolar bodies (PNBs) and subsequently re-localize to active NORs, where new nucleoli are assembled^50–52^. However, our observations suggest that nucleolar assembly during oogenesis is distinct since a large number of ec rDNA loci lack Ubtf (Fig. 5b). During early embryogenesis, nucleolar assembly uses a unique strategy. Nucleolus precursor bodies (NPBs), which contain nucleolar proteins but lack robust ribosome production activity, have been proposed to serve as scaffolds for nucleolar reactivation. However, removal of NPBs does not impair ribosome biogenesis^53^ or nucleolar formation^54^ in the developing embryo, suggesting that nucleoli can assemble *de novo* without pre-existing nucleolar material. Our data are consistent with a similar model in oocytes, in which ec rDNA activation occurs through the initiation of rRNA transcription, which subsequently recruits ribosome biogenesis factors and nucleates nucleolar assembly. Alternatively, nucleoli can form by nucleolar fission, where inactive loci may acquire transcriptional competence through the redistribution of factors from pre-existing active nucleoli. In zebrafish, stage I oocytes contain large pools of Ubtf, yet only a subset of ec rDNA loci is active (Fig 5b). By stage II, multiple ec rDNA foci frequently share Ubtf signal, and by stage III all ec rDNA foci have their own unique Ubtf signal. These observations position oogenesis as a powerful system to dissect the dynamic and stepwise assembly of nucleoli, revealing how nucleolar organization is regulated in response to developmental demand.

The canonical view of ribosome assembly is that it proceeds as a continuous and tightly regulated process^55^ that can be paused or altered in the context of cellular stress or disease^56–58^. Our findings extend this view by showing that nucleolar architecture (Fig. 2) and FC–DFC layering (Fig. 4) are dynamically regulated during zebrafish oogenesis. Because nucleolar organization is directly coupled to rRNA synthesis and processing^7^, these changes suggest that distinct steps of ribosome biogenesis are differentially regulated across developmental stages. One possibility is that oocytes temporally partition ribosome assembly, prioritizing rRNA synthesis at early stages while limiting downstream processing, and subsequently shifting toward rRNA maturation and folding at later stages. Such a relay mechanism of assembly would allow the oocyte to balance biosynthetic output with developmental needs. A conceptually similar strategy has been described in *Xenopus*, where 5S rRNA is synthesized early and stored in the cytoplasm as a 7S ribonucleoprotein particle for extended periods prior to its incorporation into ribosomes during late oogenesis^59–61^. Oogenesis proceeds through a prolonged and discontinuous developmental trajectory marked by alternating phases of growth, arrest, and reactivation, with each stage unfolding over a distinct temporal window^62–64^. In this context, a staged mode of ribosome biogenesis may be particularly advantageous, allowing oocytes to pause and resume production while preserving partially assembled intermediates. Such a “stop–and-go” strategy would enable efficient scaling of translational capacity without requiring continuous synthesis, and may provide a mechanism to rapidly mobilize ribosome assembly during key developmental transitions.

Many cells need to regulate ribosome biogenesis in response to external and internal stimuli. Exploring this regulation across diverse developmental contexts offers a powerful lens to expand our understanding of the nucleolus, not only as the canonical site of rRNA synthesis and processing, but as a dynamic and adaptable organelle that integrates genome organization, transcriptional control, and macromolecular assembly. Developmental systems such as oogenesis, early embryogenesis, and tissue differentiation expose modes of ribosome production that are difficult to capture in cell lines or unicellular organisms. These contexts reveal that ribosome biogenesis is not a uniform, steady-state process, but rather a highly regulated and plastic program tuned to the biosynthetic and physiological demands of the cell. By comparing how different organisms, cell types, and developmental stages build and remodel their nucleoli, we can uncover fundamental principles governing nucleolar function, identify rate-limiting steps in ribosome production, and gain insight into how these processes are rewired in disease states such as cancer.

**Fig. S1:**
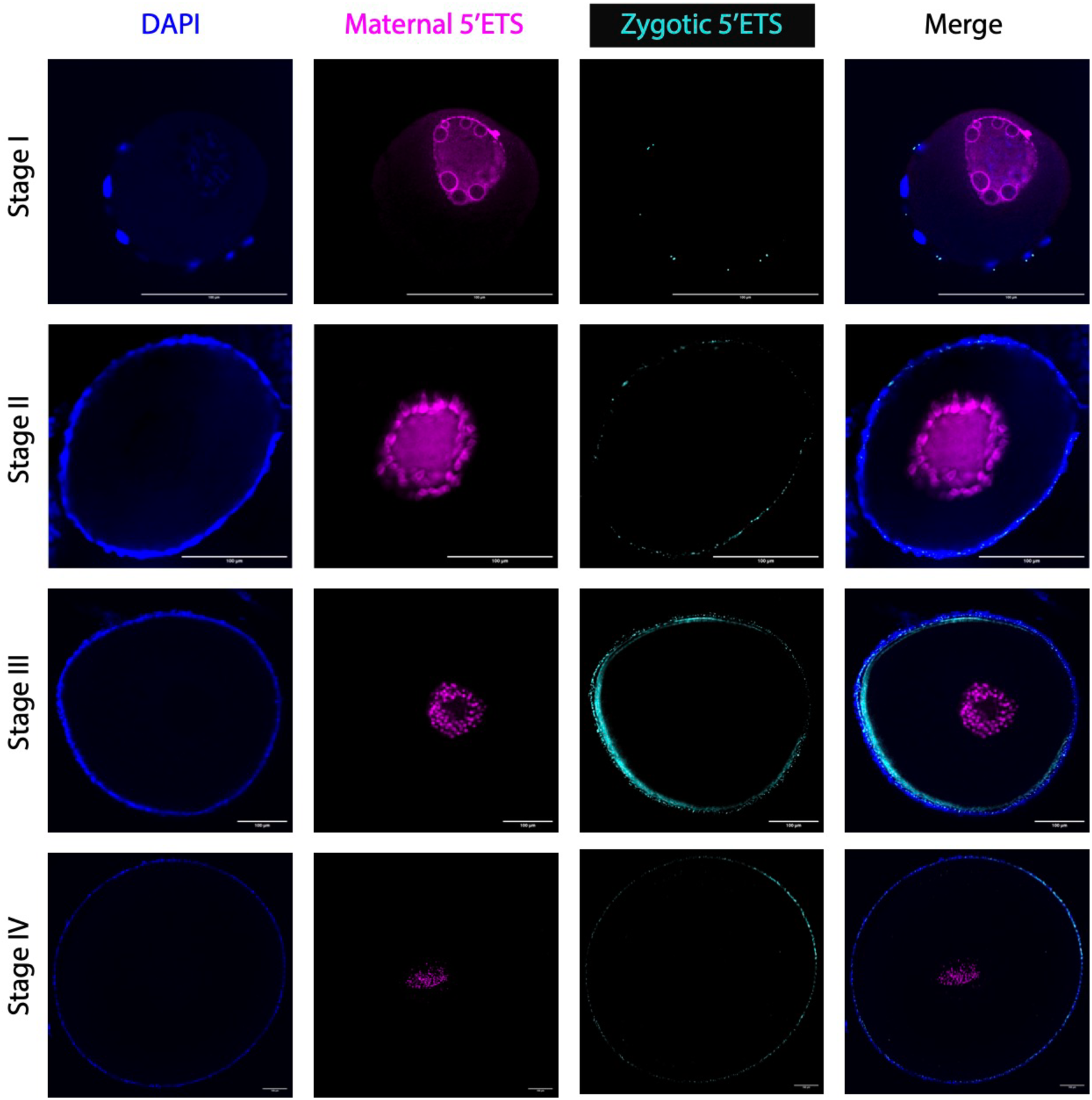
rRNA synthesis in oocytes and surrounding somatic cells. HCR RNA-FISH of nascent maternal (magenta) or zygotic (cyan) rRNA in whole-mount oocyte stages I-IV. Nuclei are stained with DAPI (blue). Note that zygotic rRNA synthesis occurs exclusively in surrounding granulosa cells, but not the developing oocyte. Scale bar: 100 µm.

**Fig. S2:**
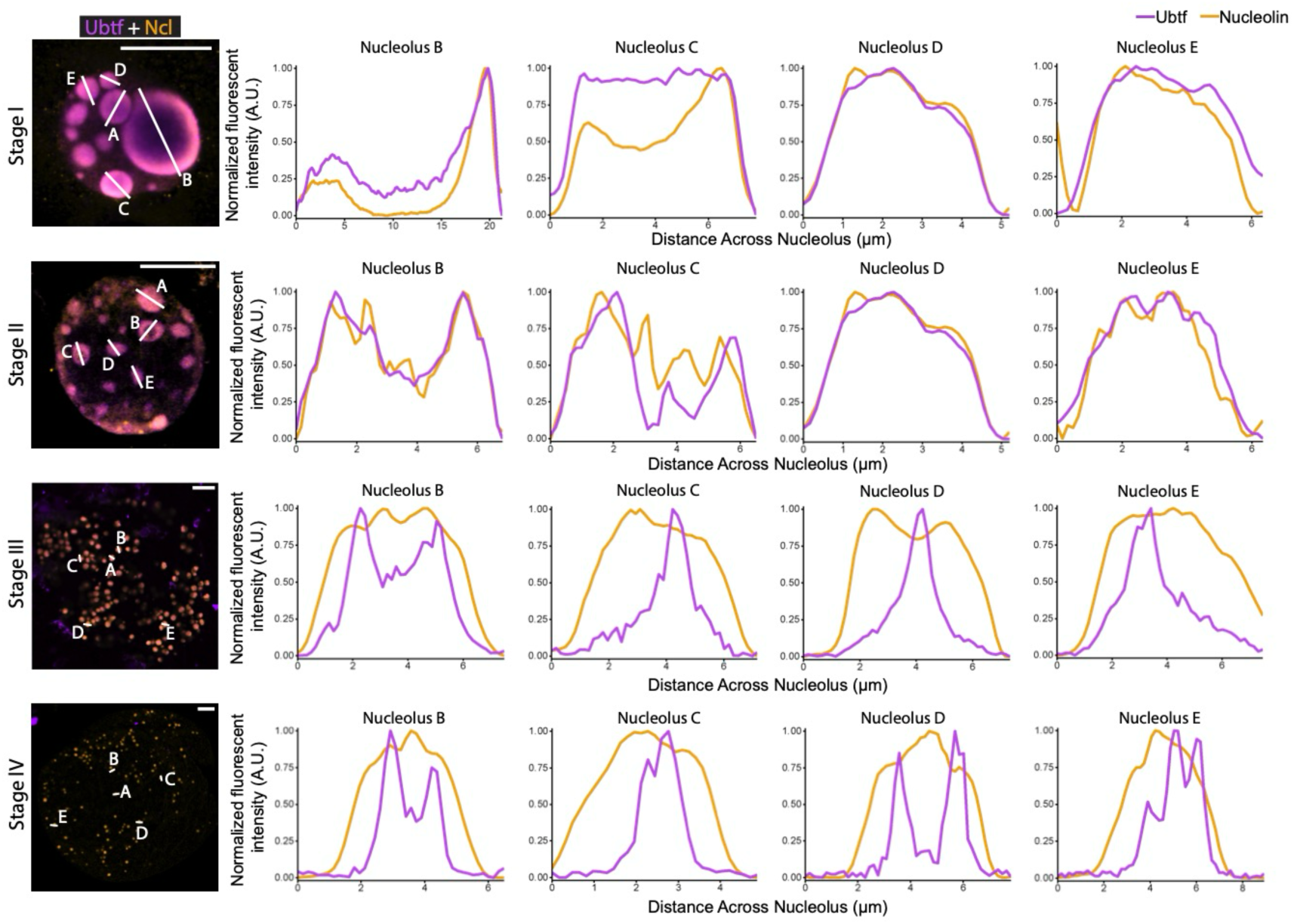
Nucleolar layers in zebrafish oocytes exhibit distinct architectures. Quantitation of fluorescent intensity measurements of nucleoli in oocytes from Fig. 4B. Oocytes were immunostained for Ubtf (purple) and Nucleolin (orange). Nucleolus measured indicated by white line and labelled A-E. Brightness and contrast were adjusted for visualization. Intensities were normalized to minimum and maximum values. A. U., Arbitrary Units. Scale bar: 20 µm.

**Fig. S3:**
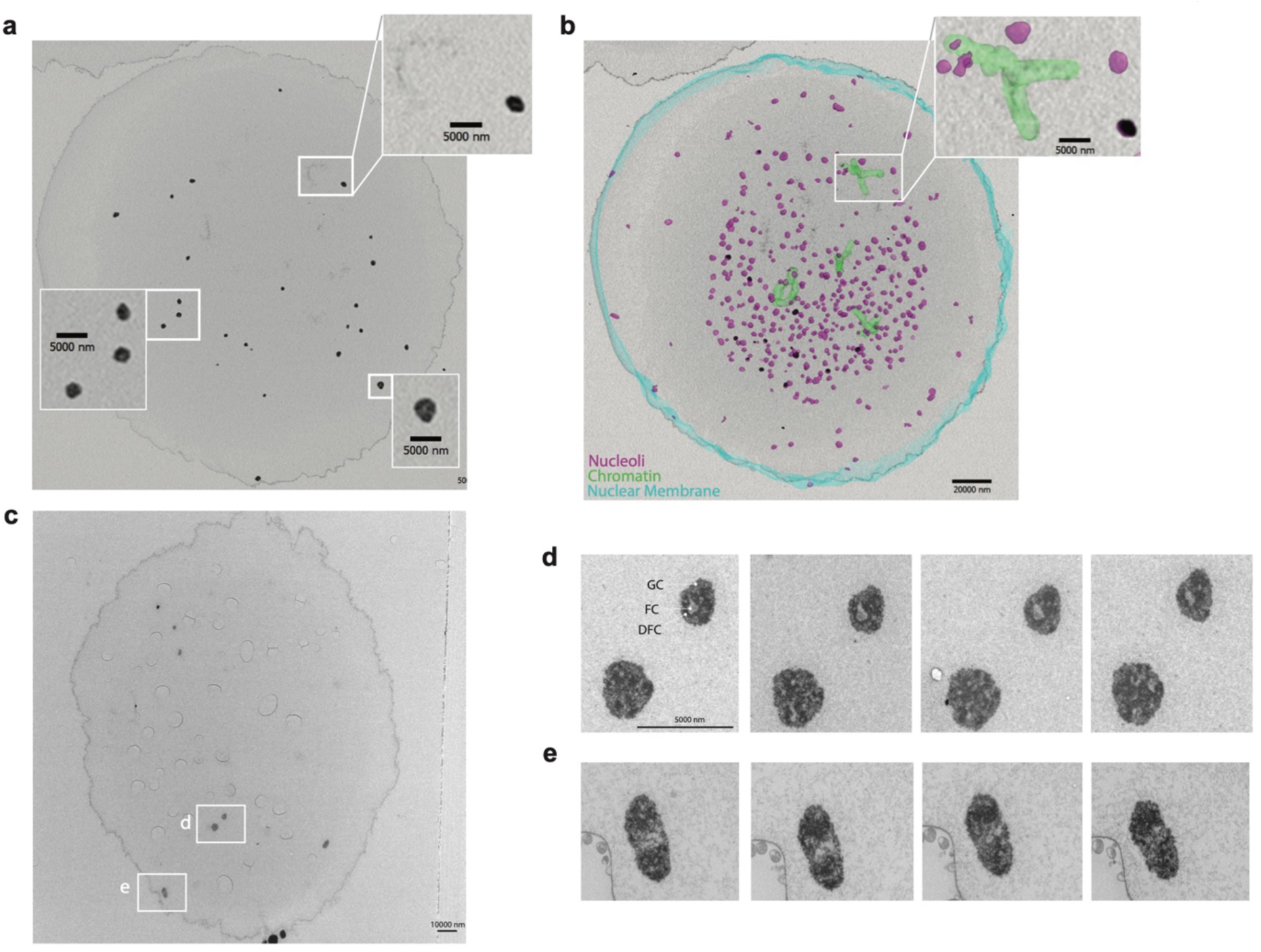
Tripartite nucleoli in stage IV zebrafish oocytes. **(a)** Array tomography of stage IV zebrafish nucleolus. Highlighted are nucleoli and condensed chromatin (white arrow). Scale bar: 5000 nm. **(b)** 3D reconstruction of nuclear structures from 89 array tomography slices, 1 µm each. Highlighted are the nuclear membrane (blue), condensed chromatin (green), and nucleoli (magenta). Scale bar: 2000 nm. **(c)** Overview of stage IV oocyte acquired with transmission electron microscopy (TEM). Scale bar: 10000 nm. Nucleoli imaged at higher resolution are highlighted. **(d, e)** Serial images through selected nucleoli showing fibrillar center (FC) (hollow centers), surrounded by dense fibrillar component (DFC), and granular component (GC). Section thickness is 100 nm, scale bar: 5000 nm.

**Fig. S4:**
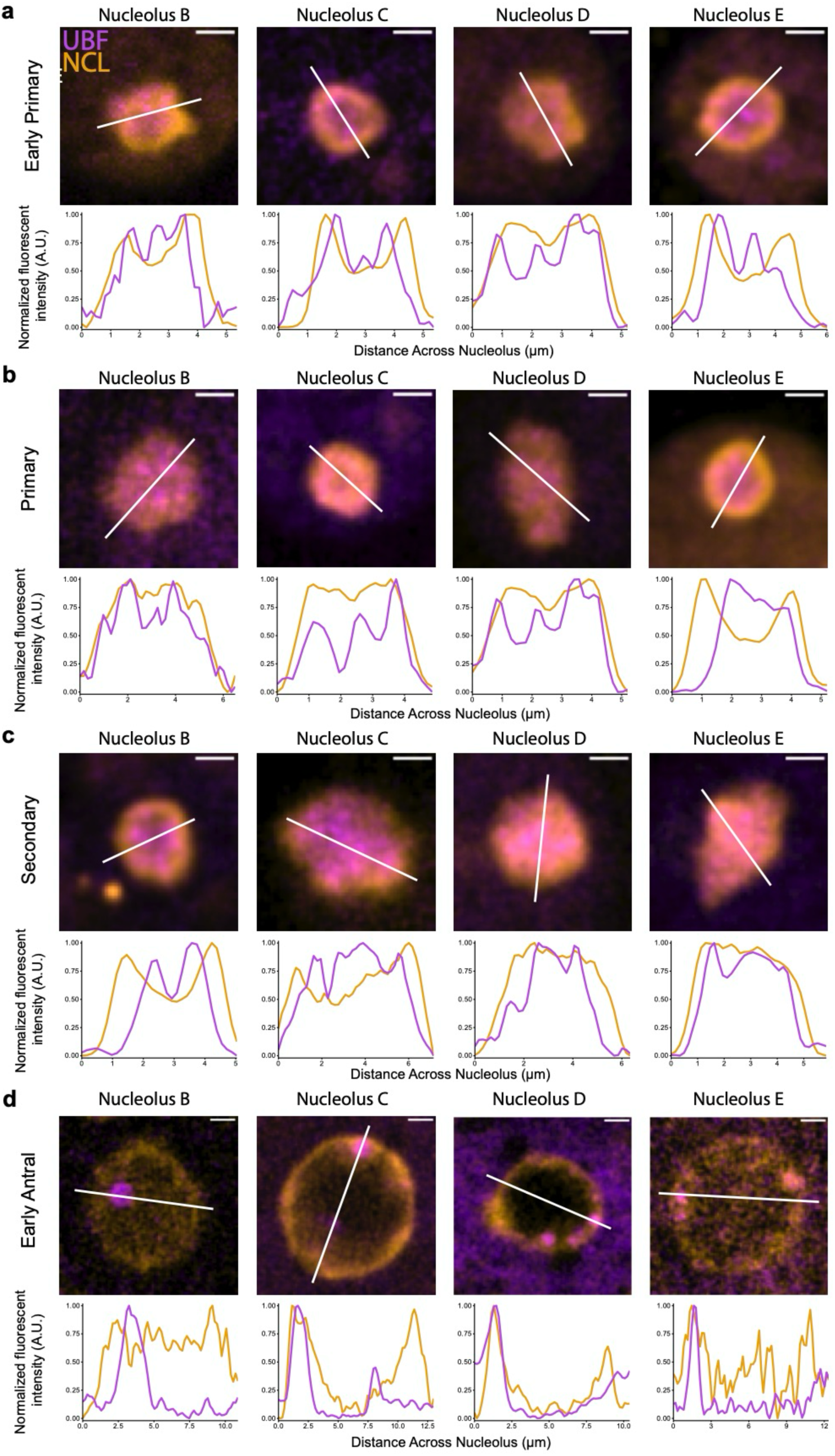
Nucleolar layers in mouse oocytes exhibit distinct architectures. **(a-d)** Quantitation of fluorescent intensities of UBF and Nucleolin across nucleoli of early primary (a), primary (b), secondary (c), and early antral (d) oocytes shown in images above. Intensities were normalized to minimum and maximum values. Early antral oocytes in panel d were imaged using a 500 ms longer exposure time for the 561 nm laser channel and 200 ms longer exposure time for the 647 nm laser channel relative to the other samples due to sample depth. A. U., Arbitrary Units. Scale bar: 2 µm.

## Acknowledgements

We thank members of the Kostova Lab for their scientific input and helpful discussions. We thank members of the Histology, Microscopy, Aquatics, and Rodents Technology Centers at the Stowers Institute for Medical Research for their support for this project. Funding: K.K. is a Freeman Hrabowski Scholar from the Howard Hughes Medical Institute. K.K. and J.L.G. are supported by funds from the Stowers Institute for Medical Research. This work was supported in part by R01HD105752-01 from NICHD to J.L.G. and F31HD116553-01 from the NICHD to A.G.

## Author Contributions

R.L., G.M., and K.K. conceived the study. R.L., G.M., D.T., C.C., and J.F.L. performed the experiments described in Fig. 1-3. S.1. G.M., D.T., T.J.C., and A.G. performed the experiments described in Fig. 4,6. S.2,4. G.M., D.T., M.M., and M.D. performed experiments described in Fig. 5. G.M., M.M., and S.H.N., performed experiments described in Fig. S3. Image quantification was performed by R.L., G.M., M.C.M., S.M., and B.R. K.K., R.L., and G.M. wrote the manuscript with input from all authors.

## Notes

https://github.com/jouyun/2026_LiMcKown

### Competing Interest Statement

The authors have declared no competing interest.

