## Supplementary material for "Nucleolar Dynamics During Oogenesis": Materials and Methods

**1. Zebrafish**

**1.1 Zebrafish husbandry and maintenance**

Adult zebrafish (AB) were kept on a 10-hour dark/14-hour light cycle at 28.5°C. All procedures comply with all relevant ethical regulations and were approved by the animal ethics committee (IACUC review board) at the Stowers Institute for Medical Research (Protocol #2024-171).

**1.2 Zebrafish dissection and ovary isolation**

Adult female zebrafish were euthanized by rapid chilling according to SOP: AQU-FSH-TEC-32. Ovaries were dissected out and immediately transferred to a Petri dish containing Leibovitz’s L-15 medium (Sigma Aldrich, L5520) or PBS (Thermofisher, J62851.AP) for subsequent processing.

For oocyte dissociation, ovaries were dissected out, rinsed in PBS, and transferred to Leibovitz’s L-15 medium (Sigma-Aldrich, L5520) supplemented with collagenase I and II (Sigma Aldrich, 17018029, 17101015) (3 mg/mL each) for stages I–II, or collagenase I alone (3 mg/mL) for stage III. Ovaries were cut into 3–4 fragments, and oocytes were dissociated by gentle pipetting with a glass Pasteur pipette and mechanical stripping with fine forceps. Stage-specific oocytes were then separated using nylon strainers (PluriSelect 43-50100-03, 43-50300-03) according to diameter, with 100 µm sieve for stage I, 300 µm sieve for stage II. Stage III oocytes were selected by checking for a central germinal vesicle (GV) under the light microscope. Stage IV were selected by checking for asymmetrically located GV under the light microscope. To minimize cell damage, dissections and isolations were completed within 1 hour. Oocytes were washed twice in enzyme-free L-15 medium before further use.

Stage V oocytes were collected by gently pressing the abdomen of gravid females anesthetized with 0.4% tricaine (Western Chemical Inc, SYNC-M-GR-US02FN) according to SOP: AQU-FSH-TEC-39. Isolated oocytes were transferred into 1.5 mL tubes in small aliquots of L-15 medium.

**2 H&E Staining of zebrafish ovary**

**2.1 Paraffin processing**

Ovaries were dissected out, rinsed in PBS, and placed in 4% PFA for fixation for 4 hours before overnight dehydration in 70% ethanol at -20°C. Samples were then placed in 100% ethanol and processed using a 3 mm paraffin embedding protocol. Briefly, samples were rinsed in 70% ethanol for 4 minutes, followed by dehydration in 100% ethanol for 10 minutes at 65 °C. Samples were then incubated in isopropyl alcohol for 45 minutes at 68 °C before paraffin infiltration through a graded series: paraffin baths 1 and 2 for 10 minutes each at 70 °C, paraffin baths 3–5 for 2 minutes each at 70 °C, and a final paraffin bath (bath 6) for 27.5 minutes at 65 °C.

**2.2 H&E staining**

Following processing, samples were embedded in paraffin wax in a sagittal orientation and sectioned on microtome (Leica HistoCore AUTOCUT - Automated Rotary Microtome149AUTOG0C1) at 4 µm section thickness onto charged glass slides (Avantik OptiBond Charged Microscope Slides SKU: SL6412-2) Slides were placed in oven at 60C for 1 hour prior to H&E Staining. Slides were stained using an H&E protocol. Slides were first incubated in an oven for 60 minutes, followed by deparaffinization in three changes of xylene (3 minutes each). Rehydration was performed through three changes of 100% ethanol (1 minute each), followed by 80% ethanol (1 minute) and a rinse in distilled water (diH₂O, 1 minute). Slides were then washed in Hemalast (30 seconds) and stained in hematoxylin for 2 minutes, followed by a 2-minute rinse in diH₂O. Differentiation was carried out for 45 seconds, followed by a 1-minute rinse in diH₂O and bluing for 1 minute. After an additional 1-minute rinse in diH₂O, slides were dehydrated in 80% ethanol (1 minute) and stained with eosin for 30 seconds. Final dehydration was performed through three changes of 100% ethanol (1 minute each), followed by clearing in three changes of xylene (1 minute each), after which slides were held in xylene until removal. Lastly, slide were coverslipped with Cytoseal XYL mounting medium (Thermo Scientific SKU: 48212-196).

**3 HCR RNA-FISH staining for whole-mount oocytes**

**3.1 Hydrogel-based clearing of zebrafish oocytes**

Dissected oocytes were fixed with 4% PFA (Electron Microscopy Sciences, 15714) for 1h, washed in PBS, dehydrated in ice-cold methanol and kept in freezer at -20C until use. Dehydrated oocytes were rehydrated through a gradient from 100% methanol to 100 mM NaHCO_3_ (Gibco, 25080-094) with 1% TritonX (Thermofisher, 327371000) in PBS and washed with 100 mM NaHCO_3_ buffer two times. Oocytes were incubated with 0.1% GMA (Glycidyl methacrylate) (Thermo Fisher #165890025) in 100mM NaHCO_3_ buffer for 1 hour at RT followed by overnight incubation at 4C, washed in PBS, and incubated in monomer solution [10% acrylamide (Thermo Fisher #J62480.AP), 0.5% N,N’methylenebisacrylamide (Thermo Fisher #L04694.22), 1xPBS] or (2.5% acrylamide, 0.6% N,N’methylenebisacrylamide, 8.625% sodium acrylate (Sigma #408220), 2M NaCl, 1xPBS) for 16 hours at 4ºC. For gelling, oocytes were incubated with fresh monomer solution for 30 min, then 0.15% tetramethylethylenediamine (TEMED) (Sigma #T22500) and 0.15% ammonium persulfate (APS) (Sigma #201531000) were added to monomer solution and incubated for 30 min. Samples were transferred to humidified chamber and incubated for 2 hours at 37ºC. After gelation, the hydrogel was incubated in clearing solution (8 U/ml proteinase K, 200 mM SDS, 200 mM NaCl, 50 mM TRIS-HCl in ddH2O) for at least 72 hours until oocytes became clear and washed in 0.1% TritonX/PBS or 0.1% Tween20/5XSSC for 5 hours.

**3.2 HCR RNA-FISH staining for** **whole-mount permeabilized oocytes**

HCR RNA-FISH was performed on whole-mount zebrafish oocytes embedded in hydrogel following Molecular Instruments’ protocol originally developed for zebrafish embryos and larvae (Molecular Instruments, MI-Protocol-RNAFISH-Zebrafish, Rev. 10^1^). Probes are listed in STable1.

**4. Microscopy**

**4.1 Confocal microscopy**

Images were acquired with an Orca Fusion 4 BT 100fps at full resolution (or Andor iXon DU897 Ultra EMCCD) on a Nikon Eclipse Ti2 microscope equipped with a Yokagawa CSU W1 10,000 rpm Spinning Disk Confocal with 50 µm pinholes. Samples were illuminated with 405 nm (3.9 mw), 488 nm (8.5 mw), 561 nm (6.1 mw), or 640 nm (46 mw) lasers (LUNV 6-line Laser Launch) with nominal power measures at the objective focal plane. This spinning disk confocal is equipped with a quad filter for excitation with 405/488/561/640. Emissions filters used to acquire images were DAPI: 430-480 nm, GFP: 507-543 nm, Red: 579-631 nm, and Far-red: 669-741nm. A Nikon Plan Apochromat Lambda LWD 40x objective lens (N.A 1.15, 0.163 um/px) objective was used to acquire the image.

**4.2 Brightfield microscopy**

Brightfield images were acquired with a Tucsen MIchrome 5 Pro color CMOS camera on a Zeiss Axiovert 200 inverted widefield microscope. Samples were illuminated with a HAL100 halogen light source and imaged using either a Zeiss Fluar 2.5x objective lens, N.A. 0.12 or a Zeiss EC Plan-Neofluar 5x objective lens, N.A. 0.16. The microscope and camera were controlled using Micromanager 1.4 acquisition software.

**5. Actin disruption**

Oocytes were treated with 4 μg/mL latrunculin A (Sigma, 428026) dissolved in 0.8% dimethyl sulphoxide (Millipore Sigma D2653) in 1xPBS for 4h at room temperature, then washed three times with 1x PBS. For subsequent phalloidin staining, stage I and stage II oocytes were immediately fixed and permeabilized after actin disruption. The nuclei of stage III and stage IV oocytes were isolated manually using fine forceps and then fixed and permeabilized.

**6. Phalloidin staining**

Oocytes were fixed in 4% PFA for 1 hour (wholemount oocytes) or 30 minutes (isolated nuclei) at room temperature, followed by three washes with 1× PBS. Samples were then permeabilized in 0.1% Triton X-100 in PBS for 30 minutes at room temperature and washed three times with PBS. Samples were incubated in Alexa Fluor Phalloidin-647 (diluted 1:400 in PBS, Invitrogen, A22287) for 1 hour avoiding light exposure at room temperature, followed by three washes of 5 minutes each in PBS.

**7. Electron microscopy of stage IV oocyte isolated nuclei**

**7.1 Sample preparation**

Nuclei were isolated from stage IV oocytes as described in section 9.1. Following final wash of nuclei in PBS, nuclei were fixed in 2.5% glutaraldehyde and 2% paraformaldehyde in 50mM sodium cacodylate buffer with 1mM sodium chloride and 1% sucrose for 2 hours before being placed at 4ºC until further processing.

**7.2 Electron microscopy**

Dissected nuclei were rinsed 3 times for 10 minutes each in buffer (50 mM sodium cacodylate buffer with 1 mM sodium chloride and 1% sucrose), incubated in 1% buffered osmium tetroxide for 20 minutes, and rinsed in ultrapure water before staining with 1% aqueous uranyl acetate for 10 minutes. After rinsing in water again, the nuclei were dehydrated through a graded ethanol series for 10 minutes each at 50%, 80%, 95%, and 2 times 100%, before transitioning to 100% acetone for 20 minutes. Nuclei were then infiltrated with 50% resin (Hard Plus Resin-812, Electron Microscopy Sciences) in acetone for 1 hour, and 100% resin overnight before embedding and curing at 60ºC for 48 hours. Cured blocks were sectioned for TEM and array tomography on a Leica UC7 ultramicrotome. Sections for TEM were cut using a Diatome ultra 45˚ diamond knife at 100 nm on formvar carbon coated slot grids, and post stained for 6 minutes each in UA (4% uranyl acetate in 70% methanol) and Sato’s lead stain (PMID: 4177281) before imaging in a Thermo Scientific Talos 200kV TEM using Velox software. Serial sections for array tomography were cut using a Diatome histo Jumbo diamond knife at 1µm on a slide and post stained with 6 minutes Sato’s lead, 6 minutes UA and 6 minutes Sato’s lead again before coating with 4 nm of carbon using a Leica ACE100 coater. The slide was imaged in a Zeiss Merlin SEM at 6 kV and 700 pA with aBSD4 detector at 300 nm pixel size using Atlas 5 software. Serial images were aligned and features segmented with IMOD^2^. Exported image sequences were converted to video using Blender 5.1.

**8. In situ hybridization (DNA)**

**8.1 Ovary cryosections**

Fixed tissue was dehydrated in a 30% sucrose/PBS solution for 1 hour. Ovary samples were individually submerged in OCT to remove excess PBS/Sucrose. Tissues were moved to a cryo-mold with fresh OCT and orientated for sagittal tissue sectioning. Tissue blocks were then flash frozen at -90°C (HistoChill, Novec™ 7000). After freezing, tissue blocks acclimated to -13°C in a cryostat (Thermo, CryoStar NX70) for 30 minutes prior to sectioning. Tissue blocks were then mounted on a cutting block with OCT and sectioned at a 5° cutting angle with 30 μm section thickness. Tissue was sectioned and collected on glass slides through the entire tissue. Slides were dried in the cryostat for 45 minutes to ensure adherence, and then stored at -70°C

**8.2 Ovary cryosection and isolated nuclei Immuno-FISH**

Prepared slides were washed in PBS three times for 5 minutes each and then treated with 0.1 mg/ml RNase A in PBS for 30 minutes at 37°C. Samples were then permeabilized in 0.5% Triton X-100 in PBS for 10 minutes before being washed three times for 5 minutes each in PBST (PBS + 0.1%Triton X-100) and then two times for 5 minutes each in PBS. Slides were then incubated in 1N HCl for 5 minutes, washed two times for 5 minutes each in PBS, and then incubated in 50% deionized formamide/2X SSC solution at 4°C overnight. Plasmids containing rDNA or maternal IGS sequence were labeled with SEEBRIGHT Green 496 dUTP or SEEBRIGHT Gold 550 dUTP using a Nick translation DNA labeling system 2.0 (Enzo Life Sciences**, ENZ-GEN111-0050**). Labeled DNA probes were denatured in hybridization buffer (Empire Genomics, Hyb-Buffer) by heating to 80°C for 5 minutes then placed on ice for 2 minutes before applying to slides and sealing under a coverslip with Cytobond (SciGene, 2020-00-1). Specimen and probe were then denatured together at 80°C for 5 minutes then hybridized in a humidified chamber at 37°C for 24-48 hours. After hybridization, slides were washed in 50% formamide/2X SSC three times for 5 minutes per wash at 45°C, then in 1x SSC solution at 45°C for 5 minutes twice and at room temperature once. Slides were washed again in PBST and blocked with 5% bovine serum albumin (BSA, Jackson Immuno Research, 001-000-161) in PBST. Ubtf primary antibody (Abnova, H00007343-M01) was diluted 1:400 and anti-Mouse AlexaFluor Plus 647 secondary antibody (ThermoFisher, A32728) was diluted 1:1000 in 2.5% BSA/PBST. Specimens were incubated with primary antibody overnight, washed three times for 5 minutes, incubated with secondary antibody for several hours, and washed again three times for 5 minutes. All washes were performed with PBST. Vectashield containing DAPI (Vector Laboratories, H-1200-10) was used for mounting. Z-stack images were acquired on a spinning disk confocal microscope with 20X, 40X, and 100X objectives.

**8.3 Hydrogel-based clearing of zebrafish oocytes for DNA FISH**

Dissected oocytes were fixed with 4% PFA for 1 hour and washed in 0.5% TritonX/PBS for 1 hour. Oocytes were washed in PBS, and incubated in monomer solution [2.5% acrylamide, 0.6% N,N’methylenebisacrylamide, 8.625% sodium acrylate (Sigma #408220), 2M NaCl, 1XPBS] for 16 hours at 4C. Oocytes were incubated with fresh monomer solution for 30 min, then 0.15% tetramethylethylenediamine (Sigma #T22500) and 0.15% ammonium persulfate (Sigma #201531000) were added to monomer solution and incubated for 30 min. Samples were transferred to humidified chamber and incubated for 2 hours at 37C. After gelation, the hydrogel was incubated in clearing solution (8 U/ml proteinase K, 200 mM SDS, 200 mM NaCl, 50 mM TRIS-HCl in ddH2O) for at least 72 hours until oocytes became clear and washed in 0.1% TritonX/PBS for 5 hours.

**8.4 Hydrogel embedded oocyte FISH**

All steps were done in a 1.5ml Eppendorf tube. Pre-cleared oocytes embedded in hydrogel were incubated in hybridization buffer (Empire Genomics, Hyb-Buffer) with 0.4mg/ml RNase A overnight at 4°C. Hybridization buffer was replaced with hybridization buffer with labeled rDNA probe and incubated at 4°C overnight. Samples were denatured at 80°C for 15 minutes and then hybridized at 37°C for 48 hours. Samples were washed in 50% formamide/2X SSC two times 15 minutes per wash at 37°C and then in 2X SSC containing 1 µg/ml DAPI four times for 1 hour each. Samples were then washed with PBST for 10 minutes and then incubated in PBS containing 1 µg/ml DAPI overnight at 4°C. Samples were washed twice in PBS for 20 minutes each and placed in a glass bottom dish (Mattek). Z-stack images were acquired on a spinning disk confocal microscope with a 40X objective (stage I and II oocytes) and a confocal point-scanning microscope with a 25X objective (stage III and IV oocytes).

**9. Immunofluorescence**

**9.1 Nucleus Isolation (Stage III-IV)**

Oocytes were dissected and washed 3 times in PBS. Stage III and IV oocytes were selected based on central and peripheral GV phenotypes and placed in fresh PBS. Ultra-fine tip forceps were used to lyse the cell membrane, and the cytosol and nucleus were gently pushed out of the oocyte. Nuclei were gently pipetted to remove cytosolic lipids before being fixed in 4% PFA.

**9.2 Manual Permeabilization (Stage I-II)**

Stage I and II oocytes are too small to lyse manually with forceps without disrupting the nucleus. To effectively permeabilize the oocyte membrane and remove some cytosol, oocyte membranes were lysed manually. First, microinjection needles were pulled from glass capillary tubes (Narishige, GD-1). Then, the needle was inserted gently into the cell membrane of stage I and II oocytes before being placed into 4% PFA.

**9.3 Immunostaining**

After fixation, oocytes and nuclei were mounted onto microscopy slides in 0.02% TritonX, 0.2 M sucrose (Fisher, S5-3), 5 mM EDTA (MillporeSigma, 324506). After drying overnight, slides were washed twice in 0.1% propylene glycol (Carolina Biological Supply, 885094), 0.04% TritonX in PBS. Isolated nuclei and whole mount oocytes were permeabilized in 0.1% and 5% TritonX in PBS, respectively. Non-specific antibody binding was minimized by blocking in HCR antibody buffer (Molecular Instruments). Primary antibodies [Nucleolin (Abcam, ab22758), and Ubtf (Thermofisher, H00007343-M01)] were applied overnight in HCR antibody buffer at 1:300 dilution. After washing 3 times in PBS, secondary antibodies [Goat anti-Mouse IgG (H+L) Alexa Fluor™ 660 (Thermofisher, A21055), and Goat anti-Rabbit IgG (H+L), Alexa Fluor™ 568 (Thermofisher, A-11011)] were diluted 1:500 in HCR antibody buffer and incubated for two hours at room temperature. DAPI (Invitrogen, D1306) was applied at 5 µg/mL for 30 min. Oocytes and nuclei were washed 3 times briefly in PBS before mounting in 70% glycerol for imaging.

**10. Mouse**

**10.1 Mouse husbandry and maintenance**

All animal studies were approved by the Stowers Institute for Medical Research Institutional Animal Care and Use Committee (IACUC protocol# 2018-0045).

**10.2 Ovary dissection and follicle isolation**

Female C57BL/6J mice (Jackson Laboratories, bred in house), approximately 6-8 weeks old, were used as oocyte donors.  Mice were humanely euthanized by cervical dislocation or terminal CO2 inhalation.  Ovaries were collected from the mice by making a small lateral incision at the midline of the abdomen using surgical scissors.  The body wall (peritoneum) was then exposed by grasping the skin firmly on both sides of the abdominal incision and pulling the skin outward toward the head and tail.  The exposed peritoneum was then carefully cut open with fine forceps and scissors to reveal the entire abdominal cavity.  The viscera were gently pushed out of the way to expose the uterine horns, which should be clearly visible on both sides of the abdominal cavity.  The ovary was removed by carefully cutting or tearing the surrounding bursa and gently pulling out the ovary with fine, curved forceps.  Isolated ovaries were placed in culture media (Advanced KSOM, MilliporeSigma) for subsequent oocyte collection.  To collect oocytes at various developmental stages, excised ovaries were transferred to a Petri dish containing pre-warmed collection media. Under a stereomicroscope, the ovary was secured with fine forceps, and a 30-gauge needle was used to puncture antral follicles by gently scoring and slicing the surface of the ovary to release cumulus–oocyte complexes (COCs). The ovarian tissue was further dissociated by gentle pipetting with a 200 µL pipette. Oocytes were allowed to settle at the bottom of the culture dish and were collected using a mouth pipette.

**10.3 Immunofluorescence**

Mouse ovary tissue was fixed in 4% PFA for 2 hours at room temperature, then washed in PBST (0.1% Tween-20, 0.5% TritonX in PBS) 3 times for 30 minutes at room temperature. Primary antibodies [Nucleolin (Abcam, ab22758), and Ubtf (Thermofisher, H00007343-M01)] were diluted in HCR antibody buffer and incubated overnight at 4ºC. Next, tissue was washed 3 times for 30 minutes in PBST. Secondary antibodies [Goat anti-Mouse IgG (H+L) Alexa Fluor™ 660 (Thermofisher, A21055), and Goat anti-Rabbit IgG (H+L), Alexa Fluor™ 568 (Thermofisher, A11011)] were diluted 1:500 in HCR antibody buffer and incubated for two hours at room temperature. The tissue was washed 3 times for 30 minutes in PBST. Then, staining with DAPI was performed (5 µg/mL) for 30 minutes followed by a brief wash in PBST. Ovary tissue was mounted on slides in 70% glycerol for imaging.

**10.4 Hydrogel-based clarification and HCR RNA-FISH staining**

Dissected ovary was fixed with 4% PFA, washed in PBS, dehydrated in ice cold methanol and kept in freezer at -20ºC until use. Dehydrated ovary was rehydrated through a gradient from absolute methanol to 100 mM NaHCO3 with 1% TritonX and washed with 100 mM NaHCO3 buffer two times. Ovary was incubated with 0.1% Glycidyl methacrylate (Thermo Fisher #165890025) in 100 mM NaHCO3 buffer for 1 hour at RT followed by overnight incubation at 4ºC, washed in PBS, and incubated in monomer solution [2.5% acrylamide, 0.6% N,N’methylenebisacrylamide, 8.625% sodium acrylate (Sigma #408220), 2M NaCl, 1XPBS] for 16 hours at 4C. Ovary was incubated with fresh monomer solution for 30 min, then 0.15% tetramethylethylenediamine (Sigma #T22500) and 0.15% ammonium persulfate (Sigma #201531000) were added to monomer solution and incubated for 30 min. Samples were transferred to humidified chamber and incubated for 2 hours at 37ºC. After gelation, the hydrogel was incubated in clearing solution (8 U/ml proteinase K, 0.5% Triton-X, 200 mM NaCl, 50 mM TRIS-HCl in ddH2O) for at least 72 hours until ovary became clear and washed in PBS for 30 min.

Paraffin processing, H&E staining, and HCR RNA-FISH staining were performed as described for zebrafish ovaries.

**11. Northern blot**

**11.1 RNA isolation**

RNA was isolated using Direct-zol RNA Miniprep (Zymo, R2050), according to manufacturer’s protocol.

**11.2 Northern blot**

Total RNA (5 µg per sample) was denatured in NorthernMax™-Gly Sample Loading Dye (Invitrogen, AM8551) (1:1 v/v) at 55°C for 30 minutes and resolved on 1.2% agarose gels prepared in 1× MOPS buffer (20 mM MOPS pH 7.0, 5 mM sodium acetate, 1 mM EDTA). Electrophoresis was performed at 70 V for 4 hours, and RNA was visualized prior to transfer. Gels were incubated sequentially in 75 mM NaOH and neutralization buffer (1.5 M NaCl, 0.5 M Tris-HCl pH 7.5) for 20 minutes each.

RNA was transferred overnight to BrightStar™-Plus Positively Charged Nylon Membrane (Invitrogen AM10102) by capillary transfer in 10× SSC (1.5 M NaCl, 0.15 M sodium citrate pH 7.0). Membranes were rinsed briefly in 2XSSC, UV-crosslinked (300 mJ/cm²), and air-dried. Prehybridization was performed in ULTRAhyb-Oligo buffer (Invitrogen, AM8669) at 42°C for 10 min, followed by hybridization overnight at 42°C with 5 nM 5′-biotinylated DNA oligonucleotide probes (STable1). Membranes were washed twice for 30 minutes at 42°C in stringency wash buffer (2× SSC, 0.5% (w/v) SDS).

For detection, membranes were washed twice in wash buffer (1XPBS, 0.5% (w/v) SDS), twice in blocking buffer (1XPBS, 0.5% (w/v) SDS, 0.1% (w/v) I-Block reagent (Invitrogen, T2015)) and incubated in blocking buffer for 30 min at room temperature. The membrane was incubated in conjugating solution (10 mL blocking buffer plus 1 µl of streptavidin-alkaline phosphatase conjugate (Invitrogen, S921)) for 30 minutes at room temperature. Membrane was washed once in blocking buffer, three times in wash buffer, and twice in assay buffer (0.1 M Tris-HCl (pH 9.5), 0.1 M NaCl). Membrane was incubated with CDP-Star substrate (Invitrogen, T2146) for 5 minutes. Chemiluminescent signal was detected using a digital imaging system Odyssey® XF. All steps were performed under RNase-free conditions.

**12. Statistics and Reproducibility**

**12.1 Quantification of HCR 3D reconstruction images**

Stage I: Segmentation of stage I nucleoli was performed using StarDist^3^. The StarDist model was trained from two hand annotated examples. Eigenvectors/eigenvalues of these segmentations were calculated using inertia tensor from scikit-image and the major, median, minor axes tabulated for each object.  Nuclei from the same images were segmented by blurring and thresholding the nucleolar images, and their axes similarly computed.  All code can be found at <https://github.com/jouyun/2026_LiMcKown>

Stage II-IV: Microscopy images of HCR labeled oocytes were analyzed in Imaris (Oxford Instruments). Nucleoli were found using the surfaces tool with 0.3 µm (stage II) or 1 µm (stages III and IV) detail and background subtraction of 1.21diameter (stage II) or 5.5 seed point diameter (stage III and IV) to separate nucleoli. Care was taken with the threshold for each image to encompass the nucleoli without extending beyond the fluorescence borders. Nuclei were annotated by manually creating a surface with contour lines in z-planes no more than 25 µm apart encircling the whole volume of nucleoli.

Measurements of the number, size and center location of the nucleoli and nucleus were exported and compared in Mathematica 12.0 (Wolfram Research, Inc). The scaled distance from the nucleolus centroid to the nucleus centroid is computed as the ratio of the actual distance between the centroids to the maximal of the nucleus semiaxes. The scaled volume of the nucleolus is defined as the ratio of the nucleolus volume to the nucleus volume. Volume of both structures approximated by the volume of the ellipsoid with the semiaxes A,B,C is given by $V=\frac{4}{3}¶ABC$.

**12.2 Quantification of HCR 3D reconstruction images for actin disrupted oocytes and corresponding controls**

Segmentation of nucleoli was performed using cellpose-SAM^4^, a Vision Transformer based version of cellpose based on the Segment Anything architecture.  Images were adjusted to have 0.5 µm isotropic resolution in X,Y,Z and processed in 3D, adjusting cellprob_threshold and flow_threshold to maximize the segmentation of the nucleoli.  Intensity and size thresholding were used to filter out aberrant objects.  All processing was done in python and utilized the scikit-image library.  Code is available at <https://github.com/jouyun/2026_LiMcKown>

**12.3 Quantification of total phalloidin intensity**

Nuclear phalloidin fluorescence intensity was quantified using Imaris (Oxford Instruments). For each oocyte, a surface was manually created using the Contour function by delineating the nuclear boundary across z-slices, reconstructing the full nuclear volume in three dimensions. Total phalloidin fluorescence intensity within each nuclear surface was extracted as the integrated intensity (sum of voxel intensities) from the phalloidin channel.

**12.4 Quantification of Ubtf and Nucleolin fluorescent intensities in zebrafish**

Images were processed using ImageJ v2.16.0. For each image, a single optical slice corresponding to the approximate center of five nucleoli was selected. Background fluorescence was reduced by applying rolling-ball background subtraction with a 50-pixel radius. A region of interest was drawn across each nucleolus, and fluorescence intensity profiles were generated for plotting.

**12.5 Quantification of Ubtf and Nucleolin fluorescent intensities in mouse**

Images were processed using ImageJ v2.16.0. For each image, a single optical slice corresponding to the approximate center of a nucleolus was selected. Background fluorescence was reduced by applying rolling-ball background subtraction with a 50-pixel radius. Gaussian blur was applied to images with sigma radius of 0.75 pixels. A region of interest was drawn across each nucleolus, and fluorescence intensity profiles were generated for plotting.

1.	Molecular Instruments, Inc. MI-Protocol-RNAFISH-Zebrafish-Rev10. https://files.molecularinstruments.com/MI-Protocol-RNAFISH-Zebrafish-Rev10.pdf (2023).

2.	Kremer, J. R., Mastronarde, D. N. & McIntosh, J. R. Computer visualization of three-dimensional image data using IMOD. *J. Struct. Biol.* **116**, 71–76 (1996).

3.	Weigert, M. & Schmidt, U. Nuclei Instance Segmentation and Classification in Histopathology Images with Stardist. in *2022 IEEE International Symposium on Biomedical Imaging Challenges (ISBIC)* 1–4 (2022). doi:10.1109/ISBIC56247.2022.9854534.

4.	Pachitariu, M., Rariden, M. & Stringer, C. Cellpose-SAM: superhuman generalization for cellular segmentation. 2025.04.28.651001 Preprint at https://doi.org/10.1101/2025.04.28.651001 (2025).
